# Local structural flexibility drives oligomorphism in computationally designed protein assemblies

**DOI:** 10.1101/2023.10.18.562842

**Authors:** Alena Khmelinskaia, Neville P. Bethel, Farzad Fatehi, Aleksandar Antanasijevic, Andrew J. Borst, Szu-Hsueh Lai, Jing Yang (John) Wang, Bhoomika Basu Mallik, Marcos C. Miranda, Andrew M. Watkins, Cassandra Ogohara, Shane Caldwell, Mengyu Wu, Albert J.R. Heck, David Veesler, Andrew B. Ward, David Baker, Reidun Twarock, Neil P. King

**Affiliations:** Department of Biochemistry, University of Washington, Seattle, WA, USA; Institute for Protein Design, University of Washington, Seattle, WA, USA; Transdisciplinary Research Areas “Building Blocks of Matter and Fundamental Interactions”, University of Bonn, Bonn, Germany; Life and Medical Sciences Institute, University of Bonn, Bonn, Germany; Department of Chemistry, Ludwig Maximilian University of Munich, Munich, Germany; Department of Mathematics, University of York, York, UK; York Cross-Disciplinary Centre for Systems Analysis, University of York, York, UK; Department of Biology, University of York, York, UK; Department of Integrative, Structural and Computational Biology, The Scripps Research Institute, CA, USA; Scripps Consortium for HIV/AIDS Vaccine Development, The Scripps Research Institute, CA, USA; École polytechnique fédérale de Lausanne, Lausanne, Switzerland; Bijvoet Center for Biomolecular Research, Utrecht University, Utrecht, The Netherlands; Utrecht Institute for Pharmaceutical Sciences, Utrecht University, Utrecht, The Netherlands; Department of Chemistry, National Cheng Kung University, Tainan, Taiwan; Graduate Program in Molecular and Cellular Biology, University of Washington, Seattle, WA 98195; Prescient Design, Genentech, San Francisco, CA, USA; Howard Hughes Medical Institute, Seattle, WA, USA

## Abstract

Many naturally occurring protein assemblies have dynamic structures that allow them to perform specialized functions. For example, clathrin coats adopt a wide variety of architectures to adapt to vesicular cargos of various sizes. Although computational methods for designing novel self-assembling proteins have advanced substantially over the past decade, most existing methods focus on designing static structures with high accuracy. Here we characterize the structures of three distinct computationally designed protein assemblies that each form multiple unanticipated architectures, and identify flexibility in specific regions of the subunits of each assembly as the source of structural diversity. Cryo-EM single-particle reconstructions and native mass spectrometry showed that only two distinct architectures were observed in two of the three cases, while we obtained six cryo-EM reconstructions that likely represent a subset of the architectures present in solution in the third case. Structural modeling and molecular dynamics simulations indicated that the surprising observation of a defined range of architectures, instead of non-specific aggregation, can be explained by constrained flexibility within the building blocks. Our results suggest that deliberate use of structural flexibility as a design principle will allow exploration of previously inaccessible structural and functional space in designed protein assemblies.

## Introduction

Polyhedral protein assemblies have evolved in nature to shield macromolecules from the surrounding environment, to spatially control chemical reactions, and to display macromolecules. These sophisticated functions have inspired the scientific community to explore natural protein assemblies for biotechnological and medical applications such as drug delivery, enzyme encapsulation, and structure-based vaccine design^1–3^. However, the portfolio of natural protein assemblies is limited and often resistant to substantial modification, restricting their use in target applications.

In the last decade, protein engineering methods have been developed that allow the generation of new polyhedral protein assemblies with structural properties tailored to specific applications. In particular, computational protein design can be used to generate novel self-assembling proteins with atomic-level accuracy^4^. Symmetric protein docking followed by protein-protein interface design has been used to generate polyhedral protein assemblies across a range of sizes, from small homomeric assemblies (∼10 nm) to cage-like architectures constructed from up to four components larger than some viruses^5–12^. Powerful new machine learning-based methods for protein backbone^13–15^ and sequence^16–19^ generation promise to make the design of custom self-assembling proteins faster and easier, enabling the generation of protein-based nanomaterials of increasing sophistication by researchers in a variety of disciplines.

Since their emergence, computationally designed polyhedral protein assemblies have been customized for numerous applications, such as encapsulation of cargos of different sizes and hydrophobicities^9,20–22^ improved circulation and tissue targeting *in vivo*^20^; cellular delivery^9,23,24^; enhanced receptor-mediated signaling^14,25,26^; molecular scaffolding for structure determination^27–29^ enzymatic co-localization^30,31^; and multivalent antigen presentation^32–37^, including in multiple vaccines currently in clinical development^38,39^ or licensed for use in humans^40–42^. Although these examples showcase the practical utility of computationally designed protein assemblies, all of them used rigid building blocks that assemble with strict point group symmetry, a feature that may limit their use in certain applications. For example, the packaging capacities of such assemblies are predefined by their sizes and inner volumes, thus appropriate self-assembling protein scaffolds must be chosen or designed for each encapsulation problem.

By contrast, many naturally occurring protein assemblies involved in cargo packaging and transport are constructed from building blocks with inherent structural flexibility, enabling them to adapt to target cargos by adopting a range of architectures. For example, the capsid proteins of many viruses can assume several closely related folds that break local symmetry and form different types of protein-protein contacts to self-assemble into multiple architectures^43–47^. This polymorphism has been previously exploited to encapsulate a variety of cargos in virus-like particles, including large DNA origami nanostructures^48–50^. Clathrin provides another clear example of protein coat adaptability: flexible clathrin triskelia interact with one another to drive the formation of coats that enclose membrane vesicles of varied curvatures and thus transport cargos of varied sizes^51,52^. Indeed, clathrin coats explore a vast assembly space and are observed to form fullerene cages (polyhedra containing 12 pentagonal faces and variable numbers of hexagonal faces) spanning multiple symmetry groups.

Beyond natural protein assemblies, flexibility and the resulting polymorphism have been recurring themes in bioengineering approaches from DNA nanotechnology to protein engineering. Symmetric 5-point star DNA motifs containing central single-stranded loops have been shown to assemble into icosahedra or larger polymorphic architectures in a flexibility- and concentration-dependent manner that requires in-plane asymmetric deformations of the 5-point star tiles^53^. Similar observations were made for a pair of complementary trimeric coiled-coil peptides that self-assemble into cage-like particles when mixed^54,55^. In MS2 bacteriophage virus-like particles it has been shown that engineered changes in the interconversion of symmetric to asymmetric capsid protein dimers, arising from conformational changes in the FG-loop, shift the assembly towards larger architectures^49^. In a remarkable tour de force, a polymorphic non-viral protein capsid derived from a bacterial lumazine synthase was evolved to form a monodisperse assembly that encapsulates its own RNA and protects it from degradation^56^. Reduced porosity and increased loading capacity arose during evolution through the acquisition of structural flexibility within individual subunits that could accommodate four related but distinct folds. Finally, the pioneering approach of designing novel protein assemblies by genetically fusing the subunits of two distinct protein oligomers in predefined orientations has also been shown to result in polymorphic assemblies when semi-flexible linkers are used^57–59^. Indeed, in some cases the assemblies can be locked into a single conformation by rigidifying the linker between the two domains^60^.

In spite of the examples described above, there are currently no computational protein design methods that controllably integrate structural flexibility to achieve multiple assembly outcomes. Here we characterize three designed self-assembling proteins that form multiple unanticipated structures that significantly deviate from the intended architectures. We find that the observed architectures can be explained by constrained structural flexibility within each building block, suggesting a general route to the controllable design of oligomorphic protein assemblies.

## Results

During the course of our work designing novel protein nanoparticles, we observed that while most experimentally validated designs accurately assemble into the intended architecture, in a small number of cases the assemblies deviate from the design model. To understand the cause of these deviations, we set out to characterize in detail three such cases. KWOCAs 18 and 70 are two one-component assemblies intended to form icosahedral and octahedral architectures from trimeric building blocks, respectively^61^, while I32-10 is a two-component assembly designed to form an icosahedral nanoparticle from trimeric and dimeric building blocks^7^ (**Fig. 1a**). Dynamic light scattering (DLS) and size-exclusion chromatography (SEC) of the purified proteins suggested that in all three cases, self-assembly resulted in mostly homogeneous assemblies (**Fig. 1b,c**). However, the measured average hydrodynamic diameter was substantially larger (25, 32, and 37 nm for KWOCA 18, KWOCA 70, and I32-10, respectively) than expected from the design models (21, 13, and 25 nm, respectively). Furthermore, small angle X-ray scattering (SAXS) profiles revealed that the experimentally observed architectures possessed structural features that markedly deviate from the design models (**Fig. 1d**). Negative stain electron microscopy (nsEM) confirmed that each design assembles into well-defined, finite architectures, but that the assembly outcomes did not conform to the design models, and in the case of I32-10 were clearly heterogeneous (**Fig. 1e**). 2D class averages defined unexpected structural aspects of the resulting assemblies: square-like features and non-spherical assemblies in the averages of KWOCA 18 did not match projections calculated from the icosahedral design model; KWOCA 70 exhibited apparent five-fold symmetry that is inconsistent with the designed octahedral assembly; and averages of I32-10 contained unexpected hexagonal pores in addition to the pentagonal pores expected for the intended icosahedral symmetry. Although we have previously observed that deviations in designed protein interfaces can lead to large architectural deviations^61^, here the coexistence of features with different apparent symmetries led us to speculate that the observed architectures arise from structural flexibility within the building blocks. All three assemblies are formed from trimeric scaffolds with a similar two-domain topology: a central helical bundle that functions as a trimerization domain and a separate helical domain that contains the protein-protein interface driving assembly (**Fig. 1f, Supplementary Table 1**). In KWOCAs 18 and 70, the trimeric scaffolds were designed by helical fusion and sequence redesign between the two domains^62^, which may introduce packing defects at the junction. In I32-10, the naturally occurring trimeric scaffold (PDB ID: 1SED) contains a small interface and hinge-like loop between the two domains that closely resembles the architecture of collectins, in which flexibility between the two domains is known to be related to carbohydrate binding^63^. We thus hypothesized that flexibility within each trimeric building block drives assembly towards the observed alternative and heterogeneous conformations.

**Figure 1.**
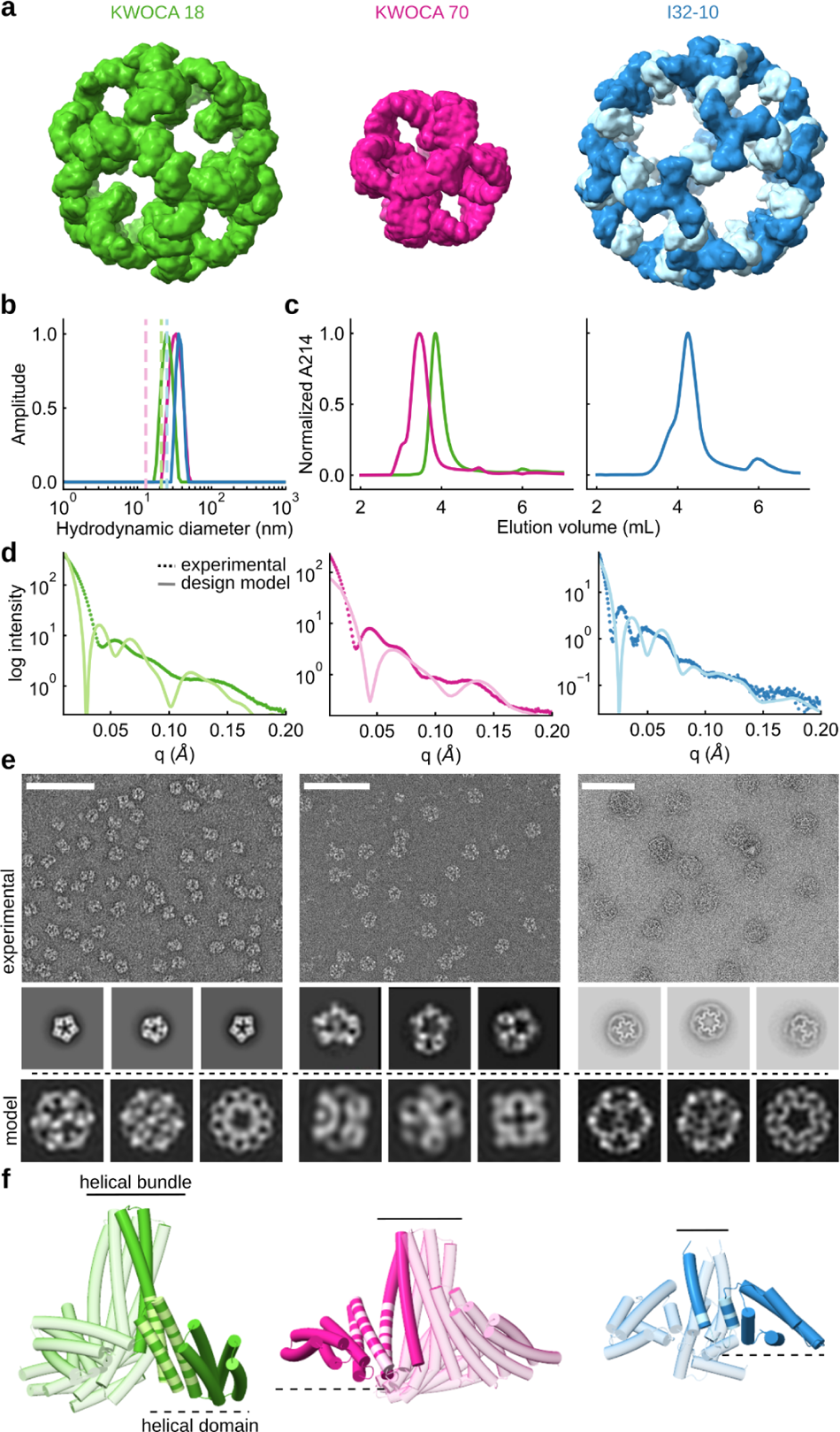
Computationally designed self-assembling proteins occasionally present large deviations from the intended architecture. (**a**) Design models of the *de novo* one-component assemblies KWOCA 18 and KWOCA 70 (ref. 61) and the two-component assembly I32-10 (ref. 7). (**b**) DLS, (**c**) SEC traces, (**d**) SAXS profiles, and (**e**) nsEM micrographs and representative 2D class averages obtained for each design. Hydrodynamic diameters, SAXS profiles, and 2D projections calculated from the computational design models are shown in light color in **b, d,** and **e**. Scale bars in **e**, 50 nm. (**f**) Representation of the two-domain structure of each trimeric building block. One subunit of each trimer is shown in a darker shade with the regions proposed to be structurally flexible highlighted in light color.

To determine whether structural metrics used during novel protein nanoparticle design could detect potential sites of flexibility, we first analyzed the interdomain region of each of the trimeric scaffolds using AlphaFold2 (AF2)^64^ and the Rosetta software suite^65^. Overall, AF2 predicted all scaffolds with high confidence (**Fig. 2a**), as assessed by the average predicted local distance difference test (pLDDT) (93, 95, and 94 for KWOCA 18, KWOCA 70, and I32-10, respectively). Previous work has correlated the degree of flexibility and the presence of unstructured regions with AF2 pLDDT at the residue level^66–68^. We thus compared in more detail the prediction accuracy within the hypothesized regions of flexibility (i.e., the junction and hinge regions) with that of core regions within each scaffold, and observed that the former present on average lower pLDDT scores. Rosetta-calculated solvent accessible surface area and average degree (i.e., the number of neighboring residues within 10 Å) additionally revealed that the junction and hinge regions appear to be more exposed to solvent and less well embedded in the protein structure than the core regions, further supporting our flexibility hypothesis (**Fig. 2b**).

**Figure 2.**
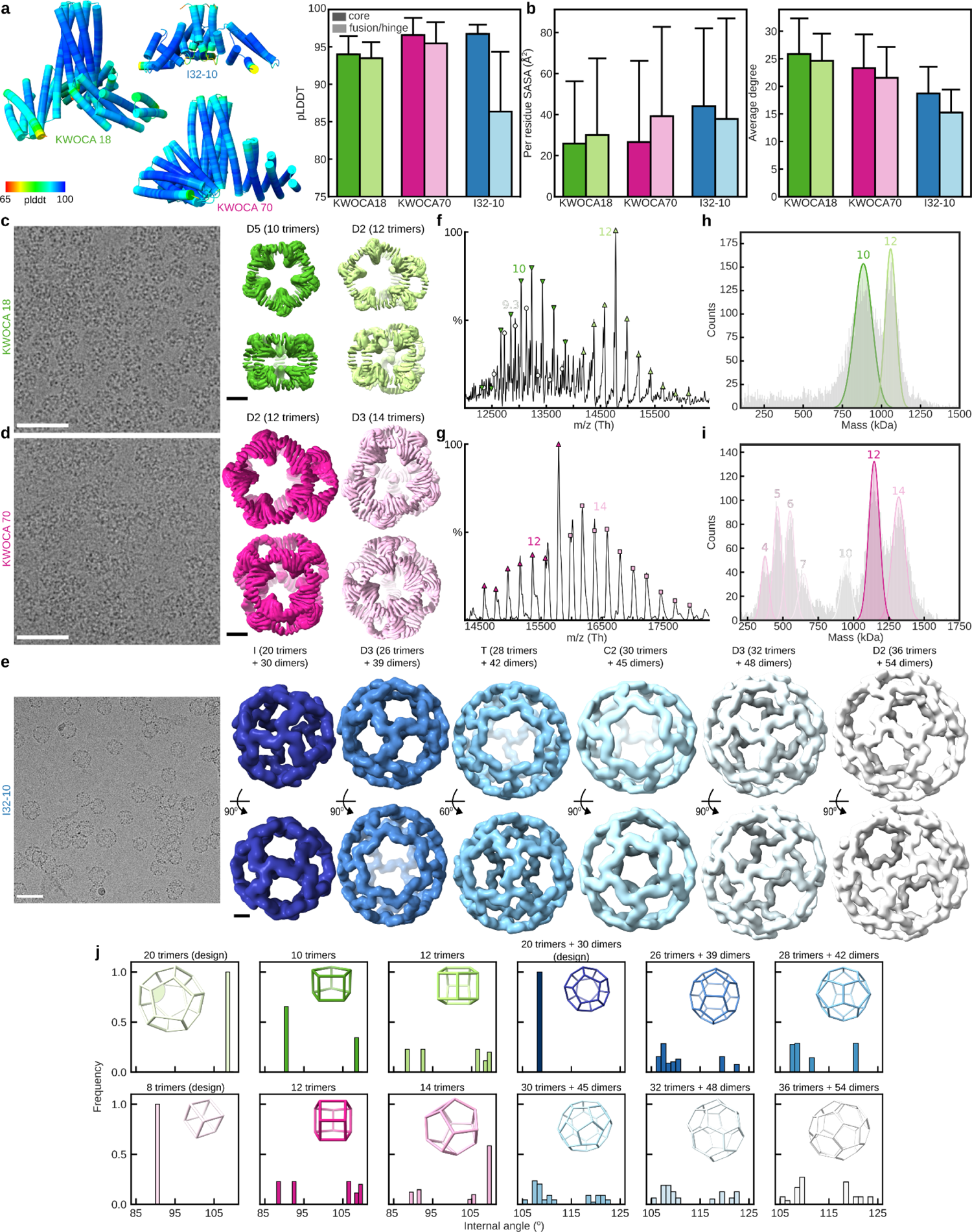
Oligomorphic assemblies of KWOCA 18, KWOCA 70, and I32-10. (**a**) AlphaFold2 multimer predictions and (**b**) Rosetta-calculated SASA and average degree suggest that the hinge regions hypothesized to be flexible are under-packed. (**c-e**) Cryo-EM micrographs and density maps obtained for KWOCA 18, KWOCA 70, and I32-10. The number of building blocks making up each assembly and its overall symmetry are indicated above each density map. Scale bars: cryo-EM micrographs, 50 nm; density maps, 5 nm. (**f,g**) native MS and (**h,i**) charge detection mass spectra obtained for KWOCAs 18 and 70. Each peak is labeled with a symbol indicating the number of trimers in the corresponding assembly species. (**j**) Distributions of internal angles in wireframe representations of the expected and observed assemblies. Wireframe assemblies are colored as in panels **c-e**, and an example internal angle is indicated for the designed icosahedral assembly of KWOCA 18.

To understand the assemblies further, we characterized them by cryo-electron microscopy (cryo-EM) (Fig. 2c-e). Micrographs of each assembly embedded in vitreous ice confirmed our assessment that they all form well-defined, finite, but apparently heterogeneous assemblies. Three-dimensional reconstructions at resolutions of 7-8 Å (**Supplementary Fig. S1, 2, Supplementary Table S2**) revealed that KWOCAs 18 and 70 do not assemble into the originally intended architectures, but instead form two distinct assemblies each, comprising both square- and pentagon-like pores: KWOCA 18 forms species with D5 and D2 symmetry assembled from 10 and 12 trimers, respectively (Fig. 2c), while KWOCA 70 forms species with D2 and D3 symmetry assembled from 12 and 14 trimers, respectively (Fig. 2d). For I32-10 we obtained six different cryo-EM maps at resolutions between 7-15 Å (**Supplementary Fig. S3**), revealing even greater heterogeneity: in addition to the originally intended icosahedral architecture (20 trimers + 30 dimers), I32-10 was also observed to form one tetrahedral (28 trimers + 42 dimers), three dihedral (26, 32, and 36 trimers + 39, 48, and 54 dimers, respectively), and one low-symmetry cyclic (30 trimers + 45 dimers) architectures all belonging to the series of fullerene cages (Fig. 2e). The internal angle distributions within wireframe models of each observed architecture highlights the flexibility required to form them (Fig. 2i). Specifically, although all internal angles are identical in the original design models due to their perfect symmetry, each of the other architectures requires a unique set of multiple internal angles. It is also important to note that in these architectures, trimeric building blocks often delineate neighboring pores of different shapes, breaking local symmetry. These observations establish that the trimeric building blocks of KWOCAs 18 and 70, and potentially the trimeric building block of I32-10, are capable of adopting multiple quasisymmetric conformations.

We then turned to native mass spectrometry^69^ (nMS) to determine whether species beyond those observed by EM were present. Consistent with the numerous assemblies observed by cryo-EM, I32-10 yielded a broad spectrum with no resolvable peaks and was not analyzed further. For KWOCA 18, three distinct assemblies containing 28, 30, and 36 subunits were readily identified in the *m/z* range between 12,000 and 16,500 (Fig. 2f). Aftercharge-state deconvolution, the acquired masses of these three architectures were 827, 886, and 1064 kDa, respectively. These measured masses are highly consistent with the calculated masses (<0.3% deviation). In the case of KWOCA 70, two different assemblies formed by 36 and 42 subunits were abundantly present in the *m/z* range between 14,000 and 18,500 (Fig. 2g). After deconvolution of charge states, masses of 1137 and 1327 kDa were obtained, respectively, both within 0.1% of the expected masses for these assembly states. Simulated mass spectra composed of three or two architectures accompanied by the calculated mass shifts fit very well the measured native mass spectra for KWOCA 18 and 70, respectively (**Supplementary Fig. S4a**). Our stoichiometry assignments were further confirmed by both Orbitrap-based single molecule charge detection mass spectrometry (CDMS) for directly acquiring mass distribution histograms (Fig. 2h,i)^70,71^ and tandem MS experiments, in which precursor ions within a certain *m/z* range were specifically selected by a quadrupole linear ion trap for the higher-energy collisional dissociation (HCD) fragmentation (**Supplementary Fig. S4b**). Both nMS spectra additionally showed a wide distribution of less abundant peaks in the lower m/z range (**Supplementary Fig. S4c**). For KWOCA 18, all oligomers from trimer to decamer could be assigned, suggesting the addition or ejection of single subunits from assembly intermediates rather than trimeric building blocks. By contrast, the number of subunits observed in smaller assembly intermediates of KWOCA 70 were all multiples of three, suggesting that the trimeric building block of KWOCA 70 is more stable than that of KWOCA 18. The relative instability and continuous distribution of assembly states may explain the presence of the 28-mer species observed for KWOCA 18. Overall these results support our observation that each design, despite its intrinsic structural flexibility, does not form disordered aggregates but instead assembles into multiple highly ordered architectures.

Intrigued by these oligomorphic assemblies, we used a combination of molecular dynamics (MD) and rigid-body fitting to cryo-EM density to investigate the origin of the flexibility. To identify inherent structural fluctuations, we performed MD simulations for each trimeric and dimeric building block as well as the interfaces formed between them (**Fig. 3a,e,i, Supplementary Fig. S5**). While the simulations suggest that the individual domains are rigid, with average RMSDs of approximately 1 Å (**Supplementary Fig. S5**), large fluctuations between trimer arms were observed. Indeed, root mean square fluctuations (RMSF) along the sequence of each monomer, which measure the average deviation of each residue from its reference position, revealed that small fluctuations along the subunit add up to large deviations at the ends of the trimer arms, the sites of the designed protein-protein interfaces that drive nanoparticle assembly (**Fig. 3b,f,j, Supplementary Table 1**). The fluctuations observed in the interface and I32-10 dimer trajectories were overall lower in magnitude than those observed in the trimer simulations. Given the observed fluctuations, we hypothesized that it should be possible to reconstruct each of the observed assemblies using a small number of structural snapshots from the MD trajectories. Indeed, one and two trimeric and two and five interface snapshots were sufficient to build well-fitting models for the cryo-EM densities of the D5 and D2 assemblies of KWOCA 18, respectively (Fig. 3c,d, Supplementary Fig. S6a). Modeling the D2 and D3 assemblies of KWOCA 70 required four and four trimeric and five and four interface snapshots, respectively (Fig. 3g,h, Supplementary Fig. S6b). Analyzing the internal angles within the snapshots used to build the models further revealed that a considerable proportion deviate from the average angles observed in the MD trajectories, highlighting that the subunits must flex to form each architecture. Notably, using rigid bodies extracted from the original, perfectly symmetric design models yielded structures that fit worse to the experimental cryo-EM densities, with large alignment defects and clashes that would prevent assembly (**Supplementary Fig. S6c-f**). Given the lower resolution of the cryo-EM maps obtained for I32-10 it was more difficult to accurately dock α-helices, so we took a simpler approach to model construction. Instead of structural snapshots from the MD trajectories, we used two building blocks extracted from the original design model, specifically chosen to introduce degrees of freedom at the main point of flexibility in the trimeric subunit: the junction/hinge region. These building blocks comprised 1) the central three-helix bundle of the trimeric component and 2) the entire dimeric component along with the two interacting globular domains of the adjacent trimeric components (**Supplementary Fig. S6a**). Rigid body fitting using these building blocks recovered all six of the observed architectures (Fig. 3l **and Supplementary Fig. S7**). Our ability to reconstruct the experimentally observed architectures for all three systems using building blocks that introduce flexibility, as opposed to perfectly symmetric rigid building blocks, strongly implicates this flexibility as the source of oligomorphic assembly.

**Figure 3.**
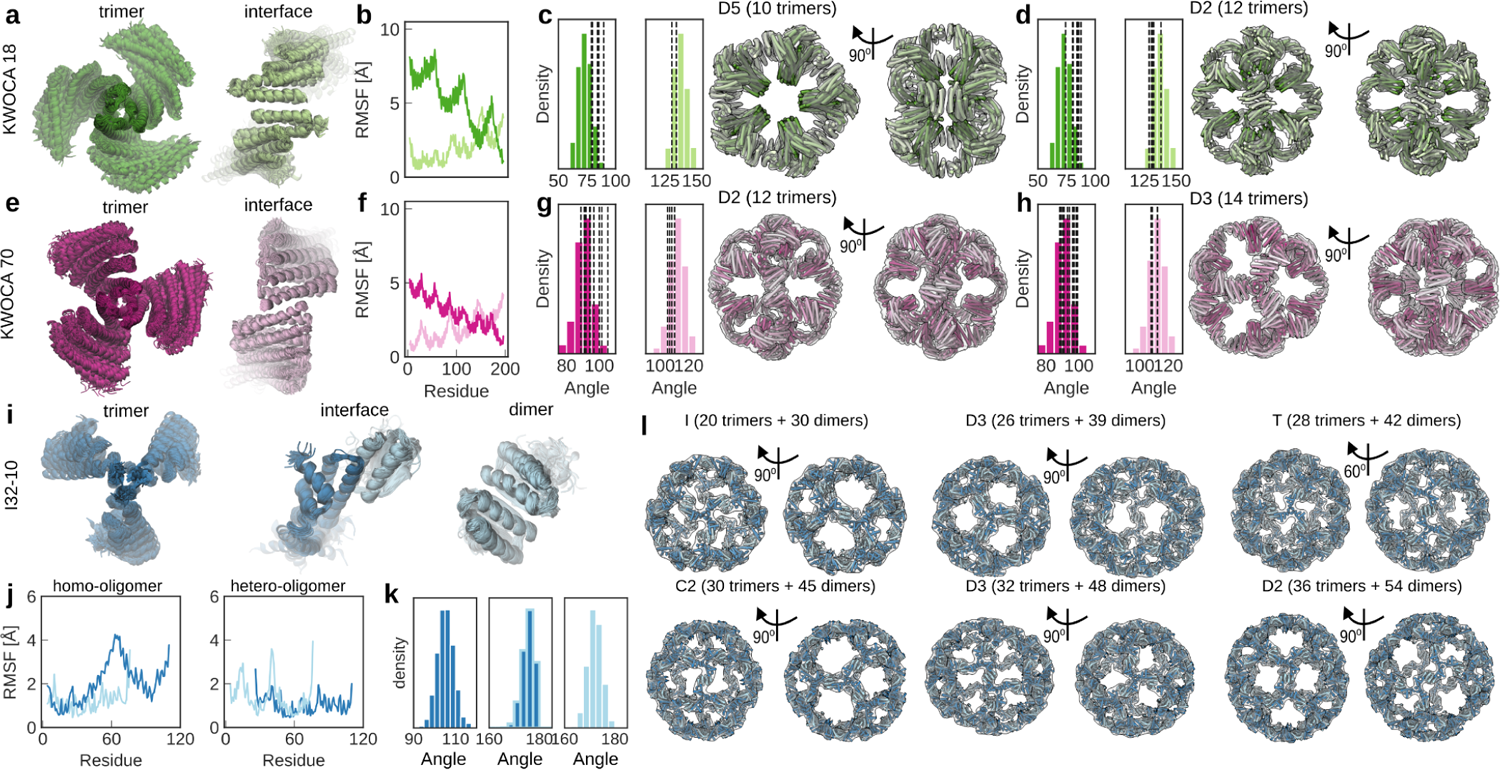
Local structural flexibility is a key driver of oligomorphism. Overlaid MD trajectories obtained for trimeric (dark) and dimeric (light) building blocks and the interfaces between them, respective root mean square fluctuations, trimer internal angle distributions, and assembly models for each observed species for (**a-d**) KWOCA 18, (**e-h**) KWOCA 70, and (**i-l**) I32-10. Internal angles within the MD trajectory snapshots used to build the KWOCA 18 and KWOCA 70 models are highlighted with dashed lines in the respective angle distributions in (**c,d,g,h**).

Given the limited number of architectures we detected by nMS and cryo-EM for KWOCAs 18 and 70, we asked what other structures might be present and at what frequency, focusing on the KWOCAs since I32-10 adopts a wider variety of architectures, likely forming even more than we were able to visualize by cryo-EM. To do so, we enumerated polyhedral architectures with regular pentagonal and square faces and evaluated their compatibility with the respective building blocks of the two systems. The key assumption behind our approach was that the geometric features of the most populated species detected by nMS and cryo-EM represent the most energetically favorable conformations of the constituent proteins. We focused on two geometric features in particular. First, we defined *S* as the sum of the internal angles *α*, *β*, and *γ* between the three subunits of each trimeric building block (Fig. 4a). *S* therefore depends on the shape of the trimer and would equal 360° if the trimer were “flat,” becoming smaller (since *α*, *β*, and *γ* become more acute) as the trimer “stretches” due to flexibility within its subunits (Fig. 4b). We calculated *S* for each trimeric building block in the KWOCA models fit into the cryo-EM reconstructions in Fig. 3c,d and found that it ranged from 278–288° in KWOCA 18 and from 305–312° in KWOCA 70 (Fig. 4c). In each case, these ranges deviated from the single values expected from the originally designed icosahedral (277°) and octahedral (318°) architectures, respectively.

**Figure 4.**
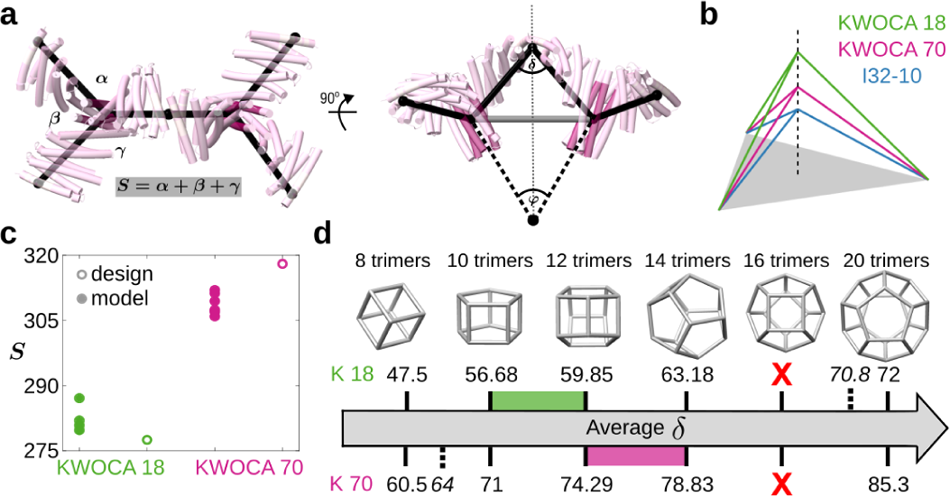
Landscapes of possible polyhedral architectures accessible to each flexible building block. (**a**) Hypothetical KWOCA architectures can be described by the sum of internal angles between the subunits of the trimer, *S* (left), and the dihedral angle *δ* at the interface between neighboring trimers (right). The internal angle between neighboring local three-fold symmetry axes, *φ*, is characteristic of each polyhedral architecture. (**b**) The trimer subunits of KWOCA 18, KWOCA 70, and I32-10 differ in *S*. (**c**) The multiple values of *S* observed experimentally for KWOCAs 18 and 70, compared to the single values of the perfectly symmetric design models, highlight the flexibility of these proteins. (**d**) The average *δ* necessary to obtain each architecture is shown. Green and pink ranges highlight the regime in the experimentally observed architectures. The 16-trimer architecture cannot be formed without extreme asymmetry (i.e., *θ* values well beyond the experimentally observed ranges). The dashed lines indicate the *δ* of the design models for KWOCA 18 (20 trimers) and KWOCA 70 (8 trimers). In both cases, *S* is outside the range of the experimentally observed architectures.

We used these experimentally observed ranges of *S* to approximate the structures of hypothetical KWOCA 18 and 70 assemblies in a series of polyhedral architectures comprising 8, 10, 12, 14, 16, and 20 trimers that includes the experimentally observed architectures for each protein (Fig. 4d). We placed “trimers” (see Materials and Methods) on each vertex of these architectures with *α*, *β*, and *γ* reflecting the angles of the underlying scaffold polyhedron. From these structures we calculated the second key geometric feature, *δ*, which represents the “interface angle” defined by the internal angle between the two “subunits” that meet at the interface between two neighboring trimeric building blocks (Fig. 4a). *δ* therefore also depends on the shape of the trimeric building block, namely *S*, as well as *φ*, the fixed internal angle between neighboring vertices in each polyhedral architecture. There is a further degree of freedom impacting *α*, *β*, and *γ* which stems from two contacting trimer arms rotating about an edge by an angle *θ* (**Supplementary Fig. S8a**), which asymmetrises the trimer angles away from those of the underlying scaffold. *θ* both affects the value of *S* and creates a twist in the concerned trimer arms. The estimated accessible range of *θ*, gauged from the cryo-EM structures of each protein, was 0–20° in KWOCA 18 and 0–12° in KWOCA 70 (**Supplementary Fig. S8b**). We posit that substantial deviations in *δ* from the values observed in the cryo-EM models in Fig. 3c,d would not be tolerated, as this would require distortion of the designed interfaces that drive assembly. We therefore estimated the feasibility of realizing each architecture by comparing the averaged values of *δ* calculated across each type of vertex to those derived from the most populated architectures observed experimentally. The range of average *δ* in the experimentally observed KWOCA 18 architectures was 56–60°, while average *δ* ranged from 74–79° in the 12- and 14-trimer architectures of KWOCA 70 (Fig. 4d). However, *δ* fell outside of the observed ranges for each of the other polyhedral architectures. For KWOCA 18 the average *δ* would have to shrink by 9° or expand by 3° to access the next architectures in the series, whereas for KWOCA 70 *δ* would have to shrink by 3° to access the 10-trimer architecture and no angle was found to satisfy the 16-trimer architecture without requiring extreme asymmetry of the trimer arms. The latter would only be feasible if *θ* increased by at least 119% for KWOCA 18 and 148% for KWOCA 70 above the maximal value in the observed range. Combined with our experimental data, these results suggest that flexibility of the trimeric building blocks and the designed interfaces between them is constrained, leading to the formation of a small number of distinct, well-defined architectures rather than a wide variety of polymorphic assemblies.

## Discussion

Here we describe how inherent structural flexibility within protein building blocks leads to the formation of a small number of well-defined architectures in computationally designed protein assemblies. While excessive flexibility is expected to result in off-target incomplete assembly or disordered aggregates^72,73^, our results suggest that constrained flexibility results in oligomorphic rather than polymorphic assembly outcomes. This has been previously demonstrated in DNA nanotechnology, where introducing flexibility in the form of short single-stranded segments within trimeric tiles enabled the generation of several distinct polyhedral architectures^53^. While current computational protein design methods mostly focus on combining symmetry with rigid building blocks to accurately design specific target structures^4^, the assemblies we describe here are quasisymmetric in that their protein building blocks have multiple related but distinct conformations. Among the best-studied quasisymmetric protein assemblies are icosahedral virus capsids, many of which are fullerene-like architectures constructed from a single capsid protein that forms pentameric and hexameric substructures consistent with icosahedral symmetry by adopting slightly different backbone conformations^74^. KWOCAs 18 and 70 and I32-10 employ quasisymmetry in a different way: the flexibility within each subunit allows for local asymmetry within the building blocks that enables the formation of assemblies with ring-like substructures of various sizes, shapes, and numbers of subunits. When compared to quasisymmetric viruses, the resulting architectures are symmetrically and morphologically quite distinct, suggesting that this design principle could be applied prospectively to explore a large space of quasisymmetric architectures previously inaccessible to design.

Such architectures and the unusual oligomorphism we observed suggest intriguing prospects for designing adaptable protein assemblies tailored to specific applications. For example, naturally occurring self-assembling systems built of intrinsically flexible scaffolds, best exemplified by clathrin, adapt to and encapsulate cargos of different sizes^51,52^. Clathrin assembles on the cytoplasmic side of intracellular vesicles, forming a coat that facilitates vesicle formation and transport. Designed proteins that form adaptable clathrin-like assemblies with defined size ranges could potentially be used as bio-orthogonal systems for encapsulating membrane-bounded vesicles, or could conceivably encapsulate other classes of biomolecules such as nucleic acids or ribonucleoprotein complexes^9,24,75^. It is intriguing to consider that interactions between such cargos and oligomorphic protein coats may shift the distribution or even type of architectures formed by the coats, allowing them to adapt to cargos of various kinds and sizes. The design principles that we have outlined here will allow this and other hypotheses to be tested, potentially leading to novel adaptable designed protein nanomaterials that can be tailored to specific applications.

## Acknowledgments

This work was funded by the Bill & Melinda Gates Foundation and the Collaboration for AIDS Vaccine Discovery (CAVD) (INV-010680 and INV-002916), the National Science Foundation (DMREF 1629214), the National Institute of Allergy and Infectious Disease (U54AI170856, 1P01AI167966, DP1AI158186, and 75N93022C00036), the Defense Threat Reduction Agency (HDTRA1-18-1-0001), Netherlands Organization for Scientific Research (NWO) (ENPPS.LIFT.019.001), the Spinoza Award (SPI.2017.028), the Wellcome Trust (Joint Investigator Award 110145 and 110146), an EPSRC Established Career Fellowship (EP/R023204/1), a Royal Society Wolfson Fellowship (RSWF/R1/180009), and generous gifts from the Audacious project and Open Philanthropy. A.A. was supported by an amfAR Mathilde Krim Fellowship in Biomedical Research (110182-69-RKVA) and N.B. was supported by an HHMI Hanna Gray Fellowship (GT11817). Structural studies at the University of Washington were supported by the University of Washington Arnold and Mabel Beckman cryoEM center and the National Institute of Health grant S10OD032290. D.V. and D.B. are Investigators of the Howard Hughes Medical Institute.

## Materials and methods

### Protein purification and low resolution characterization

Bacterial expression and purification have been performed as previously described for each nanoparticle^7,61^, with the exception of the addition of 5% glycerol to all buffers. SEC profiles for each assembly (Fig. 1) were obtained by high pressure liquid chromatography on an Agilent Bio SEC-5 500 Å (for KWOCA assemblies) or 1000 Å column (for I32-10) (Agilent) at a flow rate of 0.35 mL/min by injection of 10 μL of purified eluate using a mobile phase of Tris-buffered saline (50 mM Tris pH 8, 150 mM NaCl, 5% v/v glycerol).

Dynamic light scattering measurements (DLS) were performed using the default Sizing and Polydispersity method on the UNcle (UNchained Labs). 8.8 μL of SEC-purified elution fractions were pipetted into the provided glass cuvettes. DLS measurements were run with ten replicates at 25°C with an incubation time of 1 s; results were averaged across runs and plotted using Python. Other DLS measurements were also obtained using a DynaPro NanoStar (Wyatt) DLS setup with ten acquisitions per measurement, and three measurements per protein sample.

To identify the molecular mass of each protein, intact mass spectra were obtained via reverse-phase LC/MS on an Agilent 6230B TOF on an AdvanceBio RP-Desalting column, and subsequently deconvoluted by way of Bioconfirm using a total entropy algorithm. For LC, buffers were water with 0.1% formic acid and acetonitrile with 0.1% formic acid; the proteins were eluted using a gradient of 10% to 100% acetonitrile buffer over 2 min.

Selected SEC fractions were concentrated to 1-5 mg/mL in buffer containing 2% glycerol. The flowthrough was used as a blank for buffer subtraction during SAXS analysis. Samples were then centrifuged (13,000 g) and passed through a 0.22 µm syringe filter (Millipore). These proteins and buffer blanks were shipped to the SIBYLS High-Throughput SAXS ALS Advanced Light Source in Berkeley, California to obtain scattering data^76–79^. Scattering traces were analyzed and fit to theoretical models using the FOXS 15 server (https://modbase.compbio.ucsf.edu/foxs/)^80,81^.

For negative stain electron microscopy, samples were diluted to 0.1-0.02 mg/mL and 3 µL was negatively stained using Gilder Grids overlaid with a thin layer of carbon and 2% uranyl formate as previously described^82^. Data were collected on an Talos L120C 120 kV electron microscope equipped with a CETA camera.

### Cryo-EM

Cryo-EM grids with KWOCA 18 and 70 samples were prepared on Vitrobot mark IV (Thermo Fisher Scientific). Chamber settings were as follows: *T* = 10°C, Humidity = 100%, Blotting force = 0, Wait time = 10 s, Blotting time = 4-7s. KWOCA 18 sample was at a concentration of 4.93 mg/mL in 25 mM Tris pH 8.0, 150 mM NaCl, 2% glycerol, while KWOCA 70 was at a concentration of 4.58 mg/mL in 25 mM Tris pH 8.0, 150 mM NaCl, 5% glycerol. These samples were applied onto UltrAuFoil R 1.2/1.3 grids (Au, 300-mesh; Quantifoil Micro Tools GmbH). Prior to sample application the grids were glow-discharged using Ar/O2 plasma for 10 s on a Solarus 950 plasma cleaner (Gatan). 0.5 μL of 0.04 mM lauryl maltose neopentyl glycol (LMNG) stock solution was mixed with 3.5 μL of each KWOCA sample and 3 μl were immediately loaded onto the grids. Following the blotting step, the grids were plunge-frozen into liquid ethane cooled by liquid nitrogen.

KWOCA-containing grids were imaged on autoloader-equipped Talos Arctica transmission electron microscope (Thermo Fisher Scientific) operating at 200 kV. Raw movie micrographs were collected on K2 summit direct electron detector (Gatan) operating in counting mode. Nominal exposure magnification was 36,000 resulting in a pixel size of 1.15 Å at the specimen level. Automated data collection was performed using the Leginon package^83^. Data collection information is shown in **Supplementary Table S2**.

Raw frames were aligned and dose weighted using MotionCor2^84^ and CTF parameters were estimated using GCTF^85^. Particle picking, 2D classification and generation of initial models were performed in cryoSPARC.v3.2.0^86^. Clean particle stacks were then transferred to Relion/3.0^87^ for subsequent 3D classification and refinement steps. KWOCA 18 dataset comprised 1,803 micrographs and yielded 300,970 clean particles going into 3D. This particle subset was subjected to 3 rounds of 3D classification without symmetry imposition (C1), resulting in 2 major classes comprising 10 or 12 trimeric building blocks with apparent D5 or D2 symmetry, respectively. These classes were subjected to 3D refinement with corresponding symmetry imposed. For KWOCA 70, we collected 2,364 micrographs and obtained 360,972 particles after initial processing in cryoSPARC. Following 1 round of 3D classification without symmetry imposition (C1) we resolved 2 subsets of particles featuring 12 and 14 trimeric building blocks with apparent D2 and D3 symmetry, respectively. These classes were also subjected to 3D refinement with corresponding symmetry imposed. The 3D refined maps of different assemblies of KWOCA 18 (D5 or D2) and KWOCA 70 (D2 and D3) were post-processed (solvent masking and B-factor sharpening) and used for docking of atomic models and subsequent interpretation. Data processing workflows and relevant map statistics are displayed in **Supplementary Fig. S1**.

The I32-10 specimen was initially prepared at a concentration of 2 mg/mL and was frozen using Lacey Carbon grids, coated with a thin layer of additional carbon. To ensure even distribution of particles in ice, the sample underwent two rounds of centrifugation, first for 20 minutes and then for an additional 30 minutes in the presence of 100 mM glycine. This was done before dilution and freezing, as these steps were shown to significantly reduce the presence of aggregated and/or flocculating nanoparticles. Grids were glow discharged for 25 seconds at 15 mA, and were subsequently blotted using conditions which included a 2-2.5 second blot time, maintaining 100% humidity in the chamber, and employing 1:30 and 1:50 dilution factors with a 10-second delay before blotting. After dilution, but prior to freezing, samples were kept on ice for approximately one additional hour to allow for further equilibration of this system, which manifested as a further reduction in the amount of visible aggregation and/or flocculation of I32-10.

Cryo-electron microscopy data for the I32-10 nanoparticle were collected using a Thermo Fisher Glacios 200 kV transmission electron microscope (TEM). The microscope was equipped with a direct electron detector camera and operated at a nominal magnification of 36,000x. A total of 3,381 movies were recorded during data collection. Micrographs were acquired at a defocus range of −0.8 to −1.8 µm to ensure optimal contrast. The data collection was performed using a pixel size of 1.16 Å per pixel. Data were collected in a single-hole, one-shot-per-hole mode with stage movement for each acquisition. A total measured dose of approximately ∼75 electrons per angstrom squared (e-/Å^2) was measured during data collection.

Data processing and statistical analysis for I32-10 were conducted using a suite of computational tools and sorting methods. Initial 2D class averages and 3D ab inito models were generated utilizing CisTEM. The preliminary symmetry classification identified the predominant nanoparticle species as likely having tetrahedral symmetry, resembling a truncated triaxis tetrahedra. The data were then imported into Relion, where a series of sequential and iterative 3D classification jobs were performed, with the truncated triakis tetrahedra map used as the input reference model for the first round of 3D classification. These initial iterations revealed the additional presence of a diverse set of nanoparticle assembly configurations which deviated from the reference tetrahedra and were supported by a large amount of heterogeneity visible in both the raw CryoEM micrographs and subsequent cryoEM 2D class averages. For further 3D reconstructions, an iterative approach was adopted to parse out probable alternative assembly geometries. Initial references for 3D classifications were always derived from previous iteration outputs, focusing on intact maps of nanoparticles displaying varying symmetry groups. Notably, all 3D classification steps were executed without applying symmetry operators and in the presence of one or more initial models to help guide classification. This iterative process occasionally necessitated revisiting 2D class averages, allowing for the selective removal of particles (based on their measured 2D diameters) to help more accurately 3D classify between nanoparticle assembly states. Throughout the data processing pipeline, a comprehensive suite of software tools was harnessed, including Relion, CisTEM, and CryoSPARC. Following 3D classification and/or heterogeneous refinements within each software package, gold-standard 3D refinements were always performed, utilizing the symmetry groups identified through a visual analysis of the outputs from asymmetric 3D classification runs. The resulting refined maps were then subjected to a final round of gold-standard 3D refinement, without applying any symmetry constraints. It is noteworthy that during this final refinement stage, the initial models underwent rigorous low-pass filtering. Maps and particles that did not reach a geometry resembling the symmetrically-refined input map at this stage were promptly discarded. This decision stemmed from a reduced level of confidence due to the system’s flexibility, which led to the lower-than-expected resolution cryoEM maps reported here, and the inconsistency in achieving reproducible final map data. All high-confidence maps for I32-10 can be found here: https://files.ipd.uw.edu/pub/oligomorphism/I32-10_maps.tar.gz.

### Sample preparation for native and charge detection MS measurement

KWOCA 18, 51, and 70 samples were buffer-exchanged into 150 mM aqueous ammonium acetate solution (pH 8) using Amicon 50 kDa MWCO centrifugal filters (Merck Millipore). Then, an aliquot of ∼2 μL from each sample was directly introduced into a gold-coated borosilicate capillary (prepared in-house)for nano-electrospray ionization in positive ion mode. Eventually, all samples were analyzed on an Orbitrap Q Exactive UHMR mass spectrometer (Thermo Fisher Scientific).

### Native MS data acquisition

Instrument parameters were optimized for the transmission of high-mass ions. Therefore, ion transfer target m/z and detector optimization was set to “high m/z” mode. The in-source trapping function was enabled with a desolvation voltage of −75 V. Nitrogen was used as the collision gas with pressure ranging from 2 to 5 × 10^−10 mbar (UHV readout). Particles were desolvated in the HCD cell with HCD energies between 100 and 125 V. For tandem MS measurements, charge state distributions were isolated through a quadrupole linear ion trap, and then those subunits from selected precursor ions were ejected with relatively high HCD energies ranging from 150 to 175 V. All Data were recorded either at resolution settings corresponding to 16 or 32 ms of ion transients. A couple of scans were summed in the Xcalibur Qualbrowser to the final displayed mass spectra.

### Charge detection MS data acquisition

A resolution of 200,000 at 400 m/z was set for 1 s ion transient. The noise level parameter was fixed at 0 and the pressure level was adjusted to about 1 × 10^−10^ mbar (UHV readout). An in-source trapping voltage of −75 V was applied with optimized HCD voltage ranging from 100 to 150 V for maximal ion transmission. After the multi-scan acquisition, an appropriate calibration factor was applied which correlates the measured intensities to the charges of individual single ions as previously described^88^. According to the determined charge state, the resulting formula mass = m/z × z − z was processed to calculate the mass of each particle, separately.

### Molecular Dynamics Simulations

Each model was solvated in an octahedral periodic box of OPC water and 150 mM NaCl using AmberTools18 (ref. 89). In total, each system consisted of approximately 135,507, 131,773, 109,404, 168,225, 28,828, 93,511, 68,394 atoms for the KWOCA18 C2, KWOCA18 C3, KWOCA 70 C2, KWOCA 70 C3, I32-10 C2, I32-10 C3, and I32-10 heterodimer, respectively. Simulations were run at constant pressure (1 bar) and temperature (298 K) using the Monte Carlo barostat, the Langevin thermostat and the ff19SB forcefield^90^. Using the CUDA enabled version of Amber18, four parallel simulations for each model were equilibrated using the AmberMDprep protocol^91^. Once equilibrated, the simulations were run at 2 fs timestep for a total of 240 ns each, yielding an aggregate simulation time of 960 ns for each model.

### Model reconstruction for observed architectures

To generate reconstructed assemblies for KWOCA 18 and KWOCA 70 from the MD simulations, snapshots at 2 ns timesteps were fitted into the cryoEM maps using the *colores* function in Situs^92^. For each map, each unique trimer and dimer interface was masked and used for fitting across all snapshots. Snapshots were scored by cross-correlation, and the top 100 fits were used for stitching. The selected snapshots were then attempted for full reconstruction of the cage and adjacent trimer and dimers were scored by RMSD of an 20-residue helical region between the trimer and the assembly interface. Overall cages were then scored by overall RMSD of all stitched regions, and the cages with the lowest overall RMSD were selected for each unique nanocage. The code used for the reconstruction can be found here: https://github.com/nbethel/KWOCA_stitching.git

To reconstruct each observed I32-10 architecture, we took a different approach to introduce flexibility in the system. Two building blocks extracted from the original design model were consequently fitted into the cryoEM maps. First, the entire dimeric component along with the two interacting globular domains of the adjacent trimeric components were fitted into the maps using the *colores* function in Situs^92^. Next, unique fits not overlapping with the vertices of the maps and correctly oriented were used to position the central three-helix bundles of the trimeric component, by minimizing the RMSD between the overlapping loop region between the two building blocks that were then stitched together.

All constructed models were finally relaxed, guided by the cryo-EM densities using Rosetta^93,94^. The relaxed models can be found at: https://files.ipd.uw.edu/pub/oligomorphism/relaxed_models.tar.gz

### Classification of polyhedral scaffolds for KWOCA cages

The scaffolds of KWOCA assemblies are polyhedra formed from regular pentagonal and square faces. They can be classified using Euler’s formula. Denoting by *V*, *E*, and *F* the numbers of vertices, edges and faces in any given convex polyhedron, respectively, then *V* - *E* + *F* = 2. Let *F*_4_ and *F*_5_ be the number of squares and pentagons on the surface of the cage. Thus, *V* = (4*F*_4_ + 5*F*_5_)/3, *E* = (4*F*_4_ + 5*F*_5_)/2 and *F* = *F*_4_ + *F*_5_. Thus 2*F*_4_ + *F*_5_ = 12, implying the following solutions:

1. *F*_4_ = 6, *F*_5_= 0: cube made of 8 trimers.
2. *F*_4_ = 5, *F*_5_= 2: an assembly with D5 symmetry made of 10 trimers.
3. *F*_4_ = 4, *F*_5_= 4: an assembly with D2 symmetry made of 12 trimers.
4. *F*_4_ = 3, *F*_5_= 6: an assembly with D3 symmetry made of 14 trimers.
5. *F*_4_ = 2, *F*_5_= 8: an assembly with D4 symmetry made of 16 trimers.
6. *F*_4_ = 1, *F*_5_= 10: It is not possible to form an assembly with these values.
7. *F*_4_ = 0, *F*_5_= 12: a dodecahedron (icosahedral symmetry) made of 20 trimers.

All possible architectures are shown in Fig. 4d. Architectures are rendered using the Carbon Generator (CaGe) programme.^95^

### Mathematical reconstruction of assembly architectures

Mathematical models of KWOCA assemblies were reconstructed from the polyhedral scaffolds described above. In KWOCAs models, every trimeric protein building block is represented by three lines connecting the centre of mass of its central helical bundle (Fig. 4a, shown in dark pink) with the centres of mass of the helices forming the interfaces to neighbouring timers. In order to construct these models also for the cases when pdb files of the cages are not available, the vertices representing the centres of mass of the central helical bundles are positioned on the vertices of the polyhedral scaffold such that the helical bundle, when later mapped onto the mathematical model, aligns with the axis connecting the polyhedral centre with its vertices. They are then rotated about, and radially translated along this axis, such that lines in neighbouring trimers meet on the lines bisecting the polyhedral edges at angle *δ*. We characterise the degree of flatness of the trimer in terms of the angle sum *S* = *α* + *β* + *γ*, where *α*, *β* and *γ* denote the angles between any two neighbouring lines (Fig. 4a). The larger *S*, the flatter is the trimer. To account for the flexibility of individual arms in the trimer, we introduce the angle *θ* (**Supplementary Fig. S8**). By rotating the triangle given by the arms meeting at *δ* and the corresponding polyhedral edge around that edge, we can obtain distinct decompositions of *S*.

A polyhedral scaffold is characterised by the angle *φ* between the axis connecting neighbouring polyhedral vertices with the centre of the polyhedral frame (Fig. 4a). Given values of *S* at any two neighbouring vertices, *δ* at the connecting edge is completely determined, and vice versa. Thus, by varying *δ* and *θ* we can determine the corresponding value of *S* (code available at: github.com/MathematicalComputationalVirology/NanoMaths.git).

## Supplementary Data

**Supplementary Figure S1:**
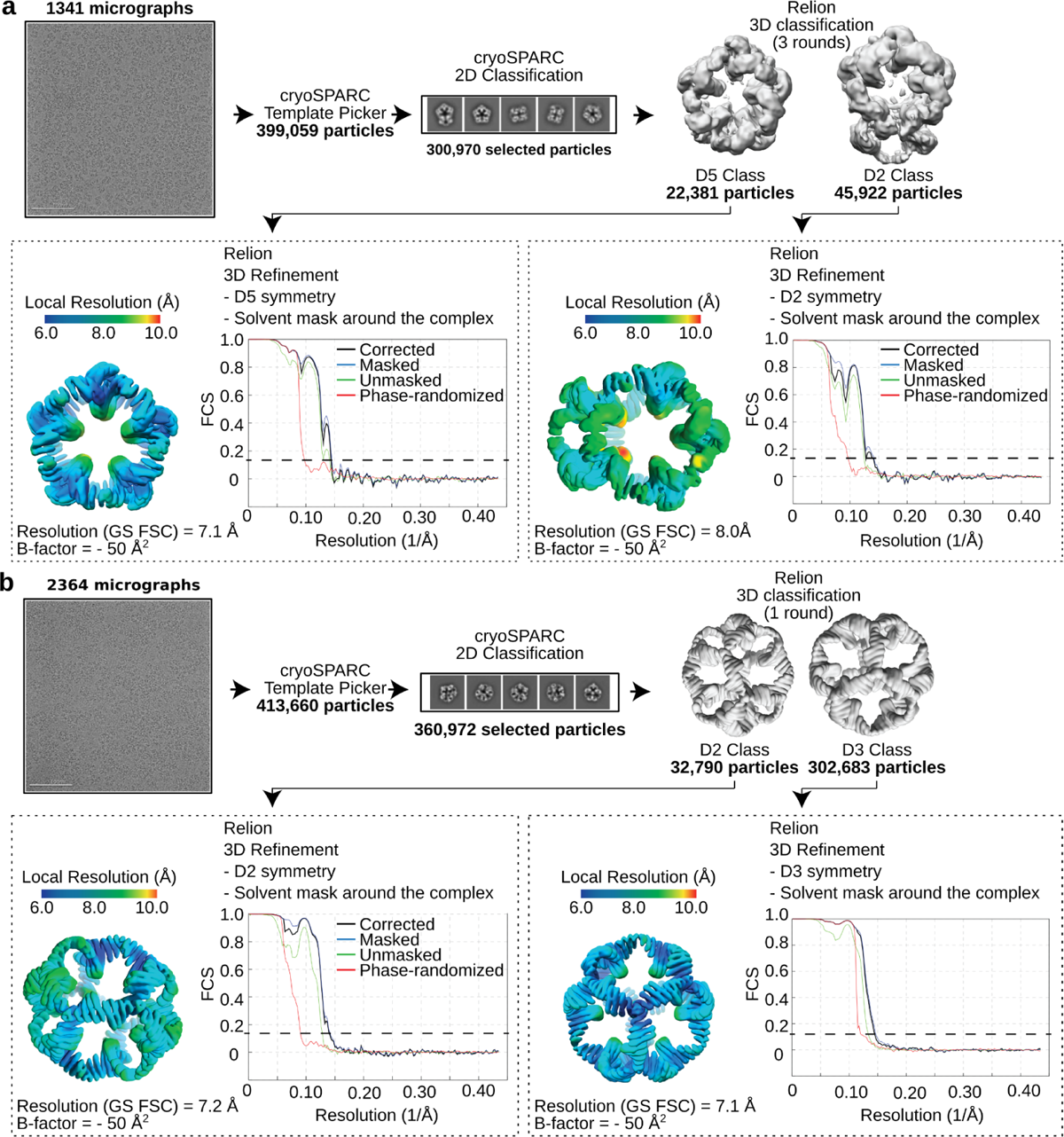
Cryo-EM acquisition and analysis pipeline for both KWOCA assemblies. Raw micrographs, representative 2D class averages, 3D classes, local resolution estimation maps, and FSC curves corresponding to the cryo-EM densities for (**a**) KWOCA 18 and (**b**) KWOCA 70 in Fig. 2.

**Supplementary Figure S2:**
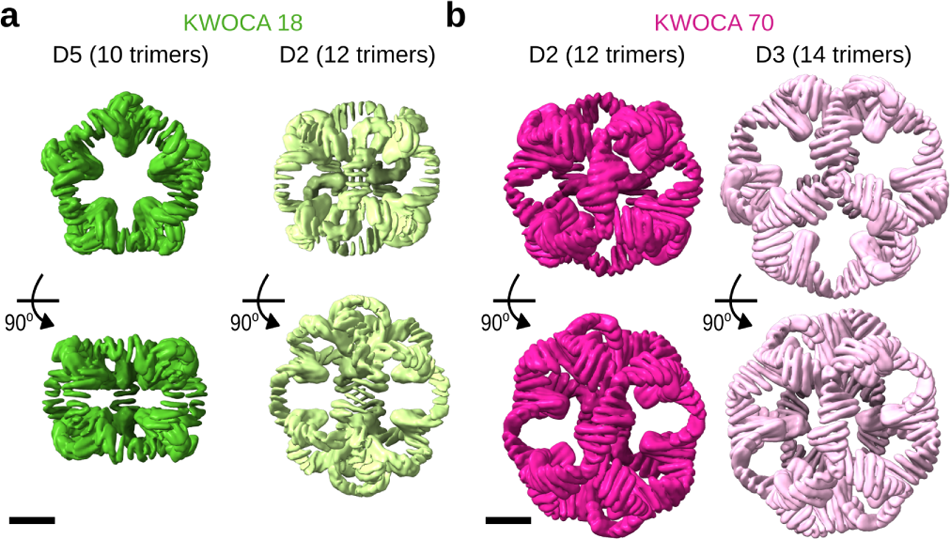
Cryo-EM density maps obtained for the KWOCA 18 and 70 assemblies. Orientations highlighting the (top) cyclic and (bottom) dihedral symmetry axes are provided in addition to those shown in Fig. 2c,d to highlight the different pores found in each architecture.

**Supplementary Figure S3:**
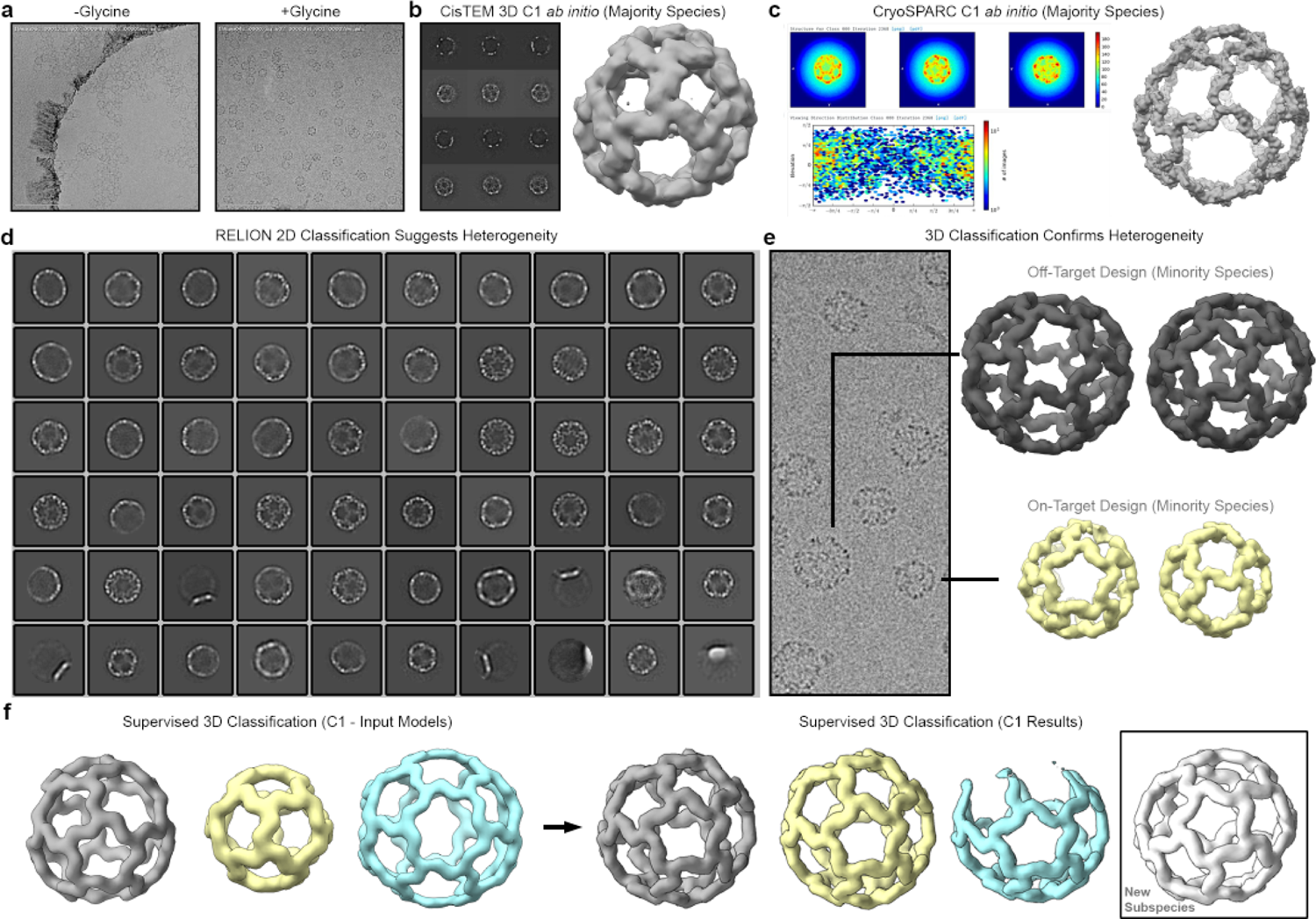
Cryo-EM insights into I32-10 assembly diversity. (**a**) Representative raw micrographs illustrate the I32-10 sample embedded within vitreous ice. The introduction of 100 mM glycine effectively mitigated flocculation/aggregation, enabling downstream data processing for the analysis of distinct I32-10 assembly states. (**b**) 3D *ab initio* reconstructions (C1 symmetry) in cisTEM unveiled a tetrahedral configuration adopted by the majority of I32-10 nanoparticles. (**c**) Independent C1 3D *ab initio* reconstructions in CryoSPARC further validated the tetrahedral assembly as the majority I32-10 species. (**d**) Relion 2D class averages highlighted remarkable heterogeneity in I32-10, encompassing variations in size and geometry. (**e**) Raw micrographs and subsequent 3D classification in Relion confirmed the existence of diverse off-target and on-target assembly states, consistent with the observations from 2D class averages. (**f**) An overview of the iterative 3D classification pipeline, as detailed in the Methods section. Supervised 3D classification jobs utilized heavily low-pass-filtered starting models, occasionally leading to the discovery of novel subspecies in output models.

**Supplementary Figure S4:**
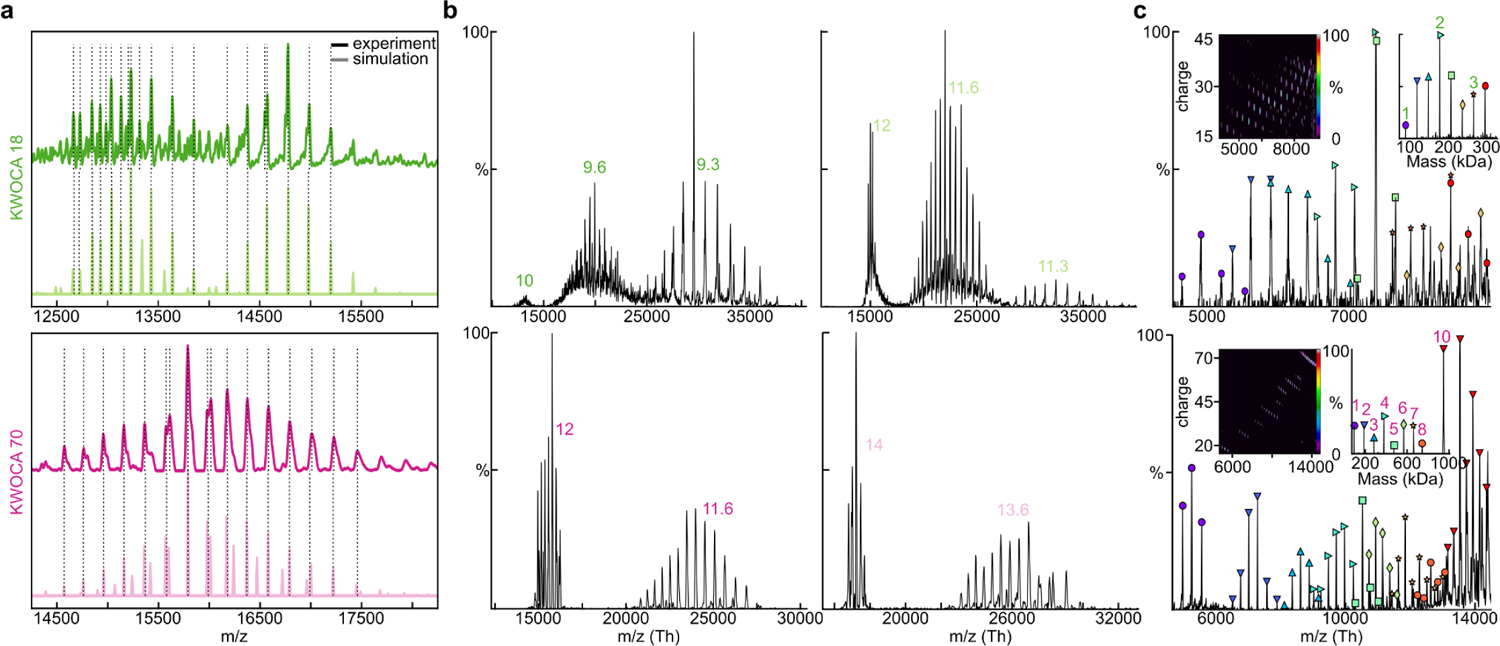
Mass spectrometry confirms coexistence of multiple species for KWOCAs 18 and 70. (**a**) Comparison between the (top) experimental and (bottom) simulated native mass spectra from Figure 2f,g. A 0.5:1:1.25 ratio of 9.3-, 10-, and 12-trimer assemblies was used in the simulation for KWOCA 18, and a 1:1.25 ratio of 12- and 14-trimer assemblies for KWOCA 70. (**b**) Tandem mass spectra by higher-energy collisional dissociation (HCD) with isolation *m/z* ranges of (left) 12,500-13,500 and (right) 14,500-15,500 for KWOCA 18, and isolation *m/z* ranges of (left) 15,000–16,000 and (right) 16,500–17,500 for KWOCA 70. (**c**) Conventional native mass spectra with lower *m/z* ranging from 4,500–10,000 and 4,000–14,500 for KWOCAs 18 and 70, respectively. The inset shows both 2D and 1D spectra of the deconvoluted masses from co-occurring low mass species.

**Supplementary Figure S5:**
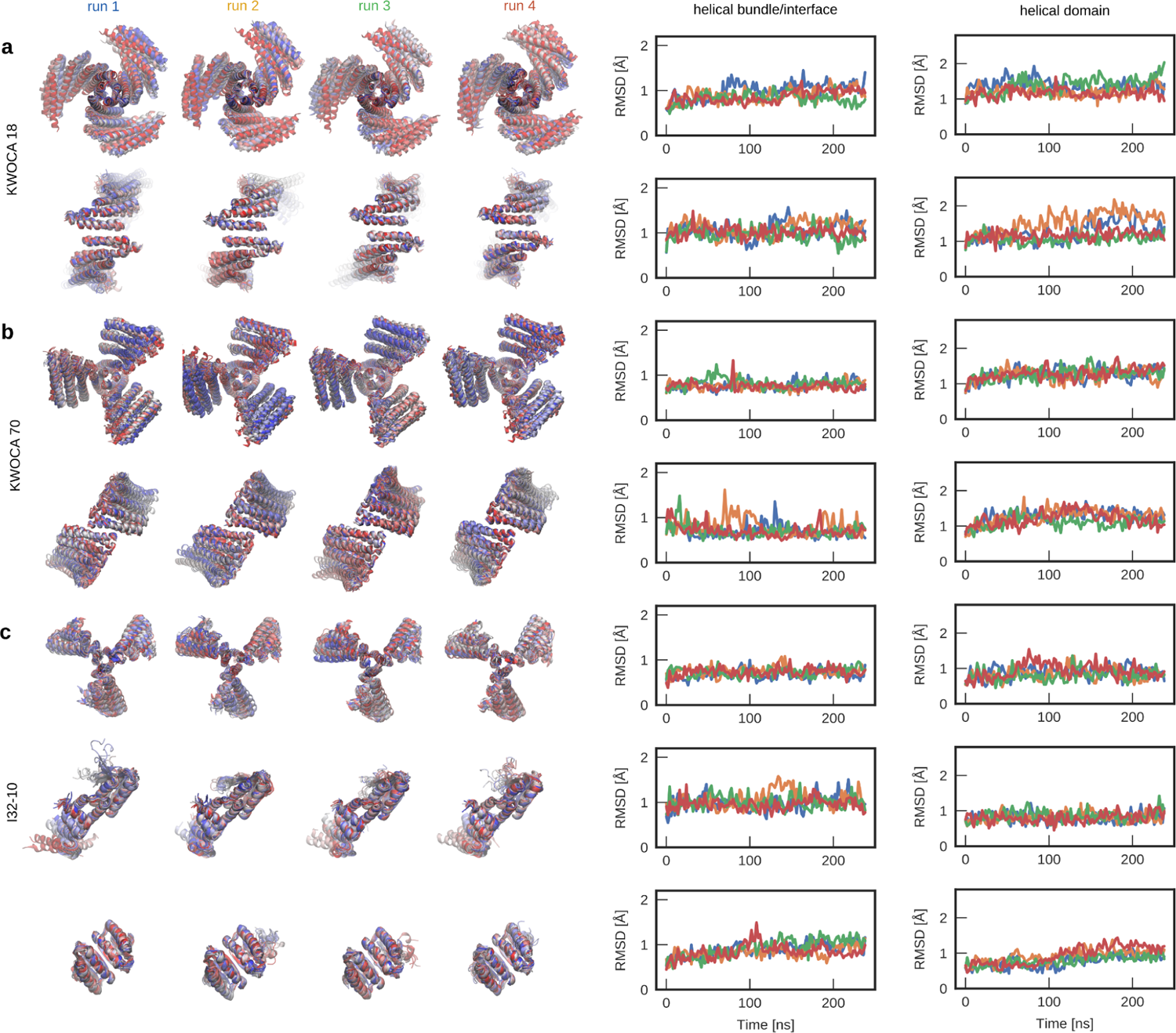
Integrity of protein building blocks, interfaces, and domains along simulated MD trajectories. Overlay of individual 240 ns trajectories (blue to red) for (top) the trimeric building block and (bottom) the interface between neighboring trimers, as well as RMSD analysis of each domain (see Fig. 1f) along the trajectories (helical bundle/interface vs. helical domain) for (**a**) KWOCA 18 and (**b**) KWOCA 70. (**c**) For I32-10, 240 ns trajectories for the (top) trimer, (middle) dimer, and (bottom) dimer-trimer interface (blue to red) and (right) RMSD analysis for each domain along the trajectories are shown.

**Supplementary Figure S6:**
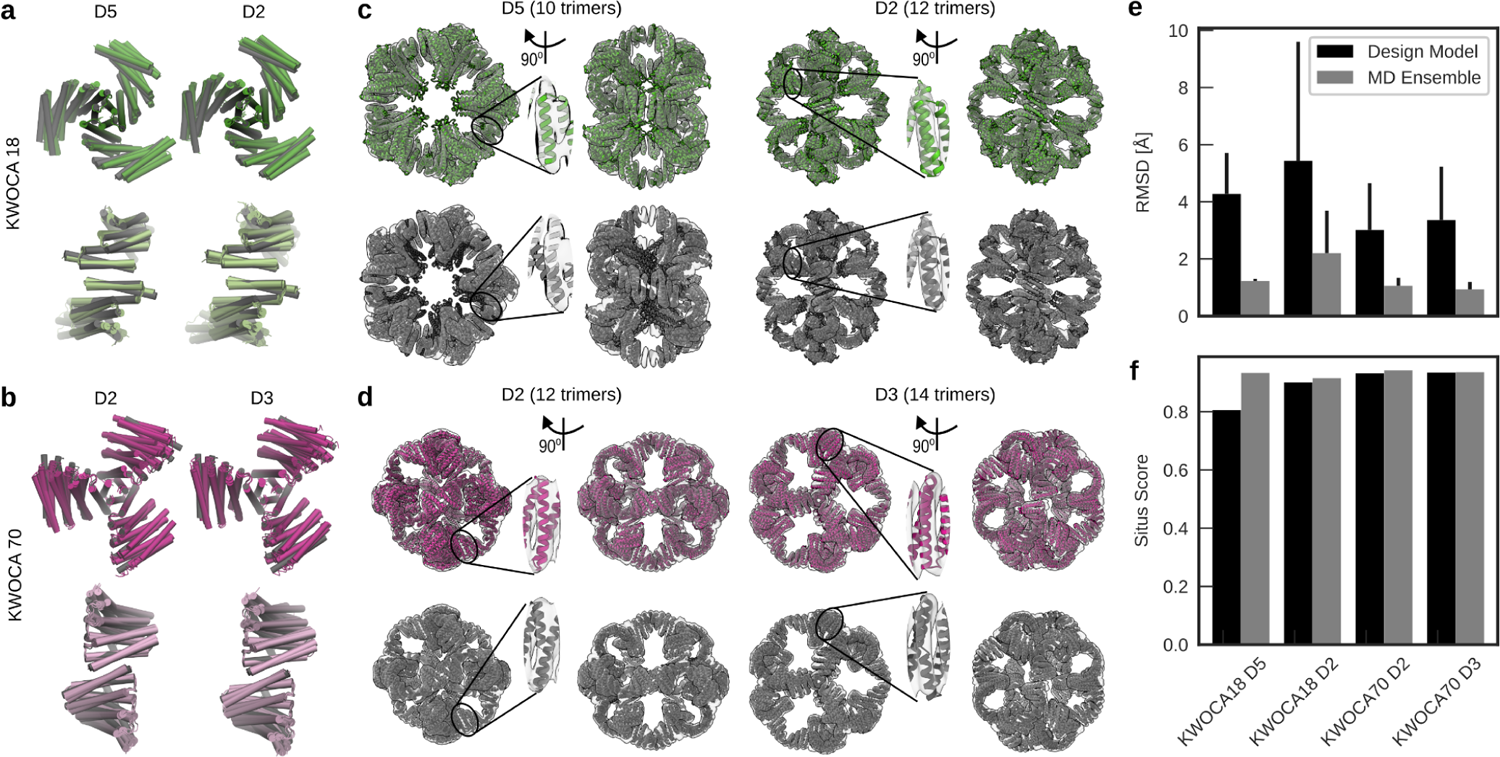
Inherent structural flexibility is sufficient to build high-quality models of the observed assemblies. (**a,b**) Overlay of the snapshots used to build each model for KWOCAs 18 and 70, respectively. Rigid bodies extracted from the design models are colored in grey. (**c,d**) Models built by fitting (top row, color) snapshots extracted from the MD trajectories or (bottom row, grey) rigid bodies extracted from the design models into the experimental cryo-EM density maps for KWOCAs 18 and 70, respectively. Insets highlight regions where the fits of the MD snapshots are clearly superior to those from the design models. (**e**) RMSDs between fitted trimeric and dimeric building blocks at the stitching region (see Materials and Methods for details) and (**f**) the overall fit of the model to the cryo-EM density using rigid bodies from the design model or snapshots from the MD trajectories both highlight the higher quality of the models constructed from the MD snapshots.

**Supplementary Figure S7:**
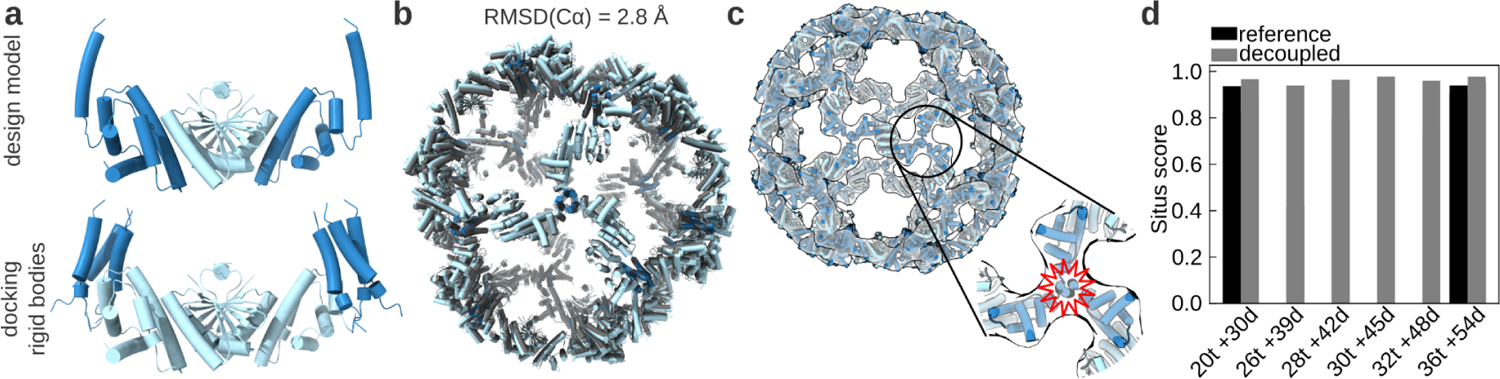
Allowing flexibility in the hinge region of the I32-10 trimer during docking renders plausible models of observed I32-10 assemblies. (a) Comparison of (top) the design model with (bottom) the rigid bodies used for two-step docking into the experimental cryo-EM density maps. In the latter, the trimeric scaffold was cropped so that the helical bundle, which occupies the vertices of each architecture (dark blue), can be treated as a separate rigid body from the helical domains that form protein-protein interfaces with the dimeric scaffold (light blue). The entire light blue region in the bottom image, comprising the dimer with both neighboring helical domains from the trimer, was used as a single rigid body during docking. (b) Overlay of the I32-10 icosahedral design model (grey) with the corresponding built model (blue). (c) Example of a fit using a single rigid body derived from the design model into the D2 architecture (36 trimers + 54 dimers) cryo-EM density. The resulting model generates clashes at the center of the helical bundle (highlighted in red). (d) Situs metric^92^ for the built models. Models for the icosahedral (20 trimers + 30 dimers) and D2 (36 trimers + 54 dimers) architectures constructed using each method are compared.

**Supplementary Figure S8:**
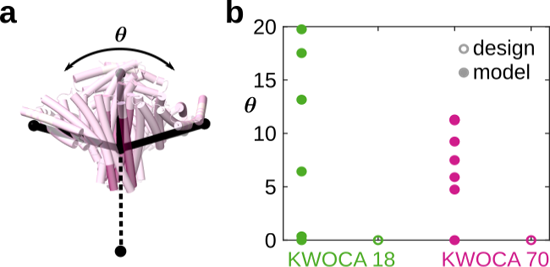
Asymmetric adjustment of individual subunits in a trimer. (a) The angle *θ* quantifies the asymmetric adjustment of the trimer. (**b**) The values of *θ* in the cryo-EM models of KWOCAs 18 and 70 compared to the designed structures. The multiple angles observed experimentally for *θ* is one indicator of the quasisymmetry observed in these systems.

**Supplementary Table S1:** List of residues comprising the junction/hinge regions of KWOCA 18, KWOCA 70, and the I32-10 trimer, as well as the core region used for comparison.

| Design | Junction/hinge region | Core region | Oligomerization interface |
| --- | --- | --- | --- |
| KWOCA 18 | 107, 108, 111, 112, 114, 115, 118, 119, 121, 122, 125, 126, 129, 134, 135, 137, 138, 141, 144, 145, 148, 149, 151-154, 156-158, 160, 161, 163-165, 167, 168, 170, 171, 174 | 54, 56, 57, 60, 61, 63-65, 67-72, 74-76, 78-81, 83, 84, 87, 88, 90, 91, 94, 97, 98, 101-103, 106, 107, 109, 110, 113, 114, 116, 117, 120, 121, 124 | 11, 14, 15, 18, 19, 21, 22, 25, 30, 33, 34, 37, 38, 40, 41, 44, 47, 48 |
| KWOCA 70 | 124, 127, 128, 131, 134, 135, 138, 145, 147, 150, 153, 154, 157, 160, 161, 164, 165, 167-174, 176-178, 180, 181, 184, 185, 187, 188, 191, 194, 195 | 30, 31, 34, 35, 37, 38, 41, 42, 44, 45, 48, 49, 51-53, 55-58, 61, 62, 64, 65, 68, 71, 72, 75, 78, 79, 94, 95, 97, 98, 101, 102, 105, 108, 109, 112 | 1-7, 9, 10, 12-17, 43, 46, 47, 50, 54 |
| I32-10 trimer | 18,19,21-27,30,33,34 | 85, 88, 89, 91, 92, 95-98, 101, 102, 104, 105, 108 | 14, 17, 21-27, 30, 32 |

**Supplementary Table S2:**
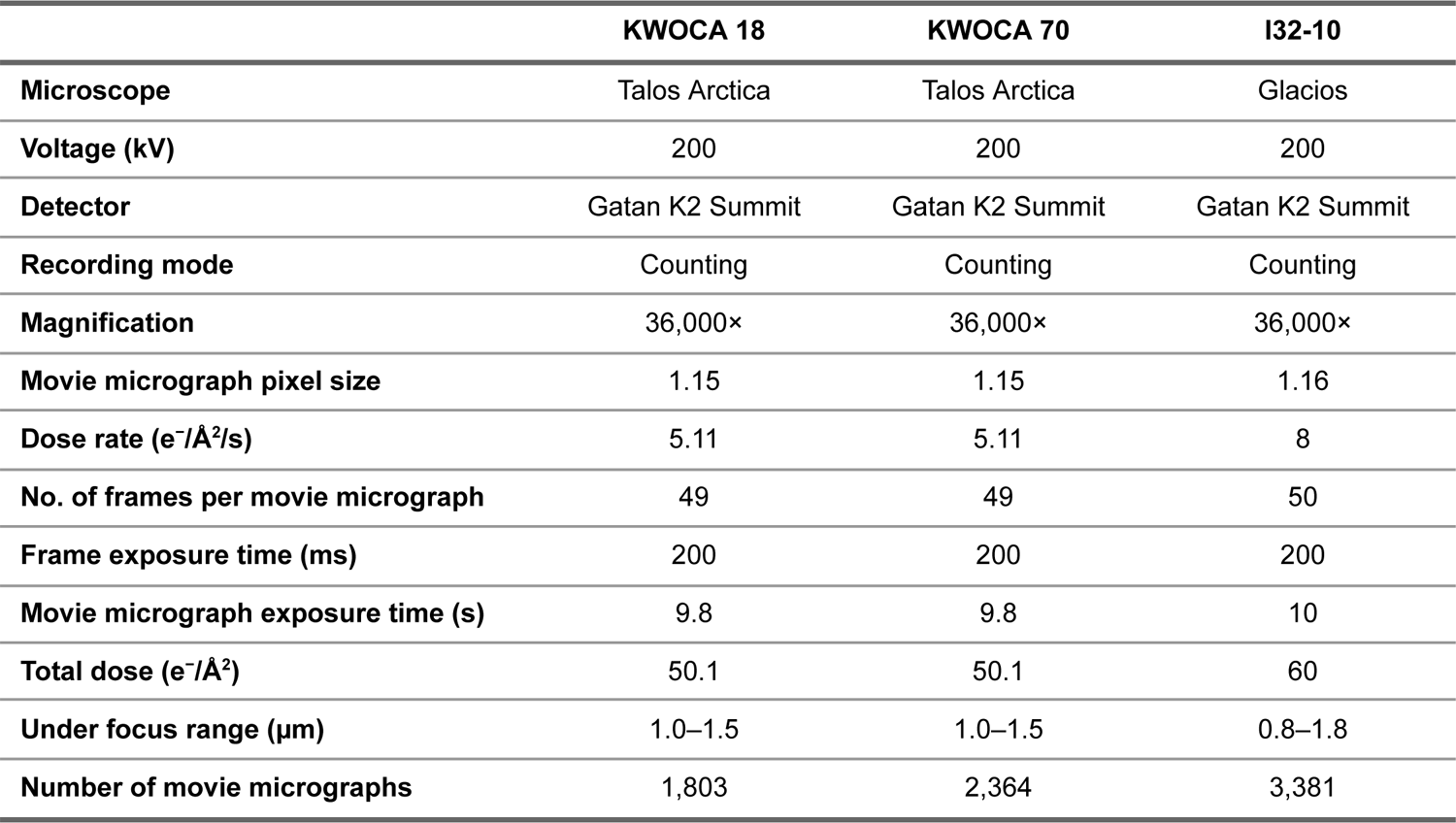
Cryo-EM data collection information.

